# Growth phase-dependent priming of galactose metabolism in *Escherichia coli*

**DOI:** 10.64898/2026.09.29.755191

**Authors:** Jordan T. Mayol, Rabia Fatima, Michael Surette, Alexander P. Hynes

## Abstract

Bacteria routinely respond to changing environments, with vastly different with nutrient availability. When nutritional changes are predictable, bacteria can deploy mechanisms to anticipate them. In these cases, an organism’s past environment can shape its response to its current environment, which is often referred to as a history-dependent behaviour (HDB). Using nutritional shift assays, we investigated how prior growth phase influences subsequent growth across different carbon sources. Here, we show evidence of a novel HDB regulating galactose metabolism in *Escherichia coli*, where previous growth to exponential phase leads to a growth advantage in a galactose environment, with no comparable effect observed in all other carbon environments tested. We demonstrate a role of the galactose repressors, GalR and GalS, in regulating this behaviour, as it is lost in strains deficient in either repressor. Further, we show how this HDB is generalizable across a collection of *E. coli* isolates, but not *Salmonella* - a closely related species with a similar natural history. Together, these findings build on the existing knowledge of HDBs governing carbohydrate metabolism by demonstrating how growth phase history can influence an organism’s response to fluctuating nutritional environments.

## INTRODUCTION

In nature, microbes are faced with a slew of environmental conditions that are subject to change, and the ability of cells to adapt to these changing conditions contributes to their remarkable success (Moreno-Gámez, 2022). Adaptation is a demanding process as it requires cells to alter their physiology to grow in a new environment. Organisms with more efficient adaptation processes often gain a fitness advantage over their competitors. For this reason, some microorganisms will use cues from their current environment to *anticipate* changes they may encounter (Mitchell et al., 2009; Schild et al., 2007; Tagkopoulos et al., 2008). In such cases, a cell’s response to a new environment can vary based on signals sensed during its past conditions. These behaviours are often related to an organism’s evolutionary (or natural) history, in which environments that are frequently encountered sequentially will select for anticipatory response strategies that function as adaptive traits (Mitchell et al., 2009). Here, we will refer to any mechanism that alters an organism’s behaviour based on its previous environment as a history-dependent behaviour (HDB). However, it should be noted that various terms have been used to describe this form of cellular memory, such as anticipatory behaviour, phenotypic memory, response memory, and hysteresis (Vermeersch et al., 2022).

Carbohydrate pathways have served as model systems for studying HDBs, with the *lac* operon being a classic example. In this system, lactose induces expression of the *lac* operon through its metabolite allolactose, which inactivates the LacI repressor and promotes production of lactose permease (LacY) and β-galactosidase (Burstein et al., 1965; Cohn & Horibata, 1959; Jobe & Bourgeois, 1972; Novick & Weiner, 1957). Since induction increases LacY-mediated lactose uptake, the system establishes a positive feedback loop that allows the induced state to persist across generations, even after lactose is removed from the medium (Maloney & Rotman, 1973). Consequently, cells with the same extracellular lactose concentration can occupy distinct, heritable expression states, demonstrating how metabolic regulation can give rise to HDBs.

The *lac* operon provides precedent for history-dependent regulation in carbohydrate metabolism and raises the question of whether similar behaviours occur in other nutritional environments. Many carbon sources work like lactose - repressed by a specific repressor, interacting with catabolite repression - a consistent model that allows for a hierarchical use of glycolytic carbon sources. In this hierarchy, glucose is preferred over lactose, and lactose over other non-glucose carbohydrates, including arabinose, xylose, sorbitol, rhamnose, and ribose (Aidelberg et al., 2014). Near the bottom of the hierarchy sits galactose, which supports substantially slower growth than other carbohydrates, reflecting the relatively low priority associated with its utilization (Tong et al., 2020). Accordingly, the *gal* regulon is tightly regulated - it is the only metabolic pathway in *E. coli* negatively regulated by two carbohydrate-specific repressor proteins, the galactose repressor (GalR) and galactose isorepressor (GalS) (Semsey et al., 2007) (Fig. 1). The two repressors share 88% amino acid similarity, inducer binding specificity, and DNA recognition sequences (Weickert & Adhya, 1992).

**Figure 1:**
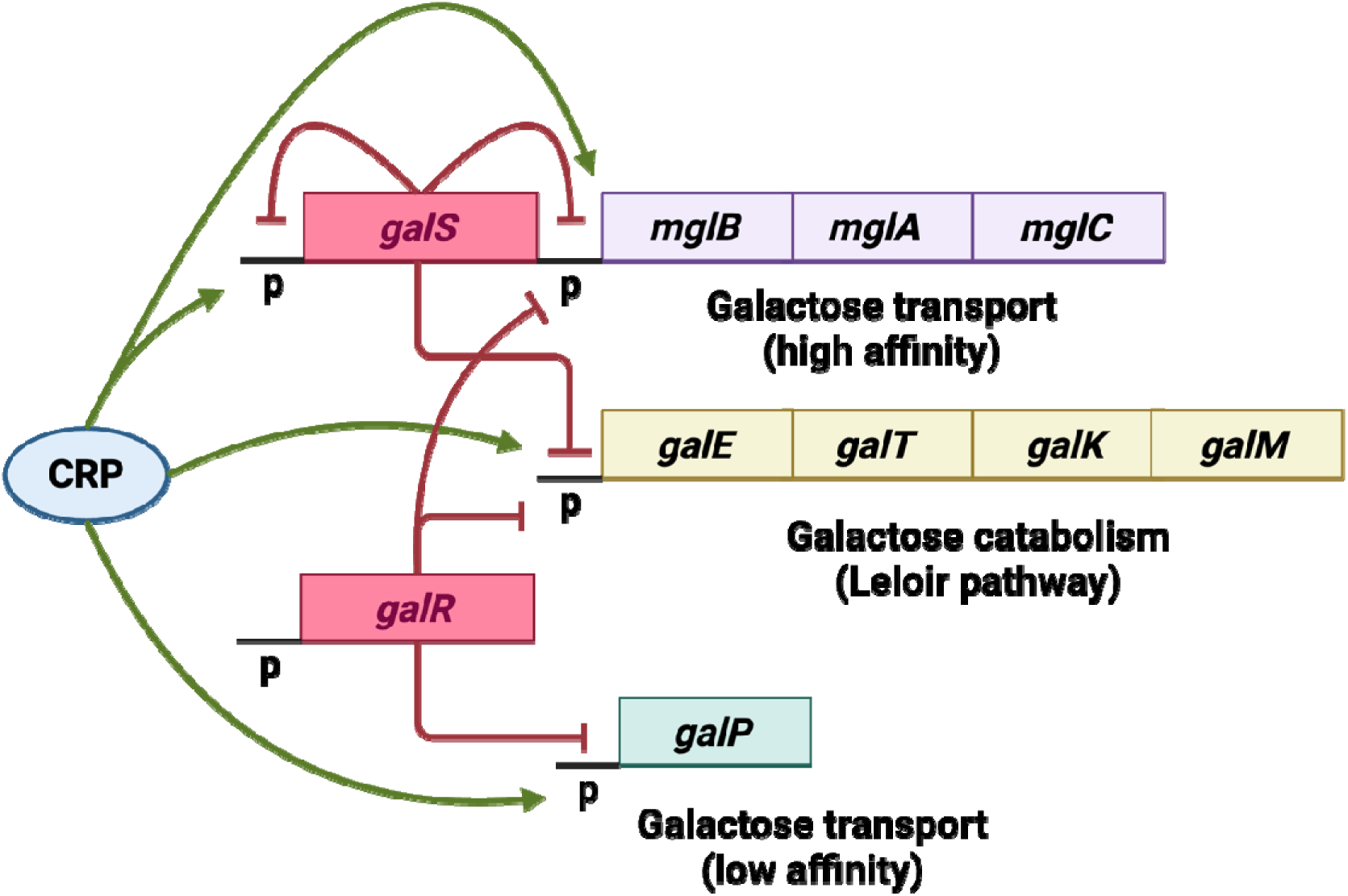
The galactose regulon of *E. coli.* Regulation of galactose transport (*mglBAC* and *galP*) and catabolism (*galETKM*) by CRP, GalS, and GalR. Green sharp arrows represent positive regulation by CRP. Red blunt arrows represent negative regulation by GalS and GalR. Promoters are indicated as “p” and coding regions are boxed. Figure adapted from Weickert & Adhya (1993).

**Figure 2:**
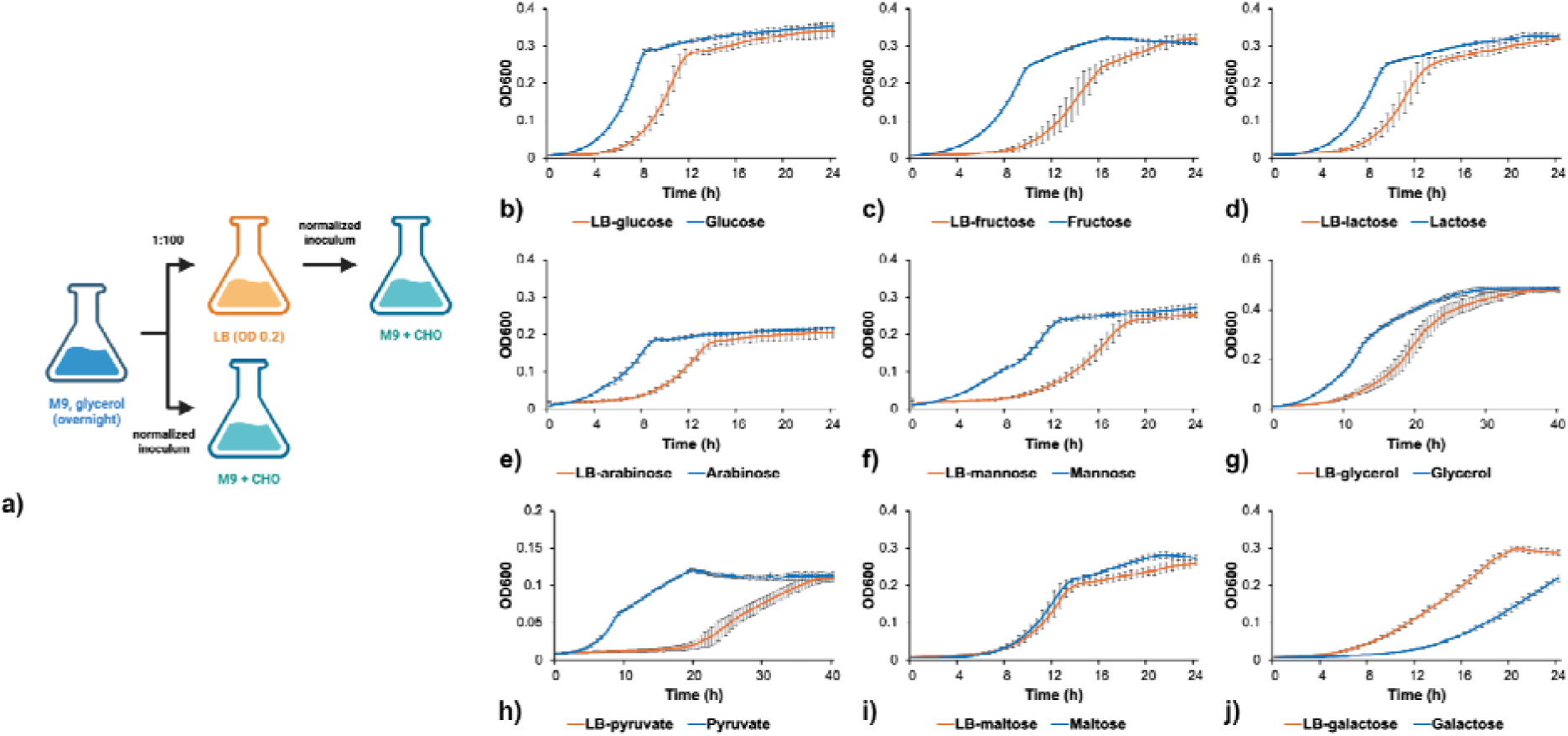
LB-to-M9 nutrient shift is disadvantageous in most carbon environments, except for galactose. (a) Diagram of LB-M9 nutrient shift assay. (b-j) Growth curves of *E. coli* MG1655 cultures previously grown to either OD_600_ 0.2 in LB (orange) or inoculated directly from M9+glycerol overnight culture diluted to the same OD_600_ (blue) and transferred to M9+galactose. Data are represented as the mean OD_600_ ± standard deviation of three technical replicates.

In the absence of galactose or its analog fucose, galactose repressors bind to operator sites in the cis-regulatory regions of promoters for genes in the *gal* regulon (Majumdar & Adhya, 1984; Weickert & Adhya, 1992). Each repressor exists as a homodimer in solution (Geanacopoulos & Adhya, 1997; Majumdar et al., 1987), and GalR can tetramerize and form higher order oligomers that enhance repression (Geanacopoulos & Adhya, 2002). Specifically, it has been shown that GalR dimers bind operator sites (*O_E_* and *O_I_*) flanking the gal operon, and DNA-bound GalR dimers can tetramerize to form a DNA loop that strengthens repression (Geanacopoulos & Adhya, 2002). In addition to negative regulation by GalR and GalS, transcription of genes in the *gal* regulon is positively controlled by the cyclic AMP receptor protein (CRP) (Choy & Adhya, 1992, 1993; Geanacopoulos & Adhya, 1997). Each of the cis-regulatory regions in the *gal* regulon contain a CRP-binding site that activates its associated promoter when glucose is scarce, due to cAMP-CRP complex formation (Weickert & Adhya, 1993).

Together, the abundance and diversity of regulatory elements controlling galactose metabolism enable the *gal* regulon to exhibit multiple levels of activity in response to the availability of galactose, fucose, and glucose. To our knowledge, these carbon sources remain the only studied stimuli influencing expression of galactose transport and catabolic genes, with glucose availability being the dominant stimulus that represses *gal* genes, even in the presence of known inducers like galactose.

Here we use nutritional shift assays to investigate history-dependent responses to different carbon sources in *E. coli*. We identify a novel HDB in which previous growth to exponential phase provides a growth advantage following transfer to a galactose environment, but is costly in other carbon environments. These findings demonstrate that history-dependent responses can emerge from the regulation of carbohydrate metabolism and provide insight into how the regulatory architecture of galactose metabolism may contribute to cellular memory.

## RESULTS

### Galactose metabolism is uniquely primed following exponential growth

*E. coli* was shifted from LB to M9 minimal media supplemented with a single carbon source to evaluate the effect of nutrient depletion on carbon metabolism. This assay was performed using a panel of carbon sources, including various carbohydrates, the tricarboxylic acid cycle intermediate pyruvate, and glycerol. Growth on each carbon source was compared to an unconditioned (LB-naïve) control culture, which was inoculated directly from an M9+glycerol overnight culture to avoid induction of pathways related to carbohydrate metabolism and minimize the potential for catabolite repression.

We found that passage through LB hindered growth in minimal media for virtually all carbon environments tested. This was expected given the huge metabolic burden for cells to shift from LB – an environment where amino acids and vitamins are in excess (Sezonov et al., 2007), back to minimal media – an environment where these essential nutrients must be synthesized *de novo*. Maltose and galactose were the only exceptions to this trend, where growth was either unchanged or improved following passage.

The ability of galactose to overcome the cost of the LB-to-M9 transition was puzzling. We hypothesized that this effect could be due to one of three things : (1) a component of LB priming galactose metabolism through induction of the *gal* operon, (2) a function of growth phase, by which cells grown to exponential phase are better equipped to handle the transition to galactose as a sole carbon source, or (3) a suppressive effect of glycerol on galactose metabolism.

A modified nutrient shift assay was performed to tease apart the effects of growth phase and LB exposure on galactose metabolism. Here, the conditioning medium (LB) was replaced with the same medium as the overnight culture (M9+glycerol) (Fig. 3a). Similarly to the LB-to-M9 nutrient shift assay, cells previously grown to exponential phase outperformed cells shifted to M9+galactose in stationary phase, demonstrated by a reduction in lag time of 2.2 ± 0.42 h (*p*<0.0001), which was used as a proxy for the relative fitness of the cultures (Fig. 3b-c). We quantified lag time using the time required for the initial population of the culture to double in optical density. Although this measure does not capture the full duration of lag phase, it provided a consistent measure that allowed us to detect and compare differences in lag between conditions. Ultimately, these findings ruled out the effect of LB on priming and demonstrated that priming was tied to exponential growth.

**Figure 3:**
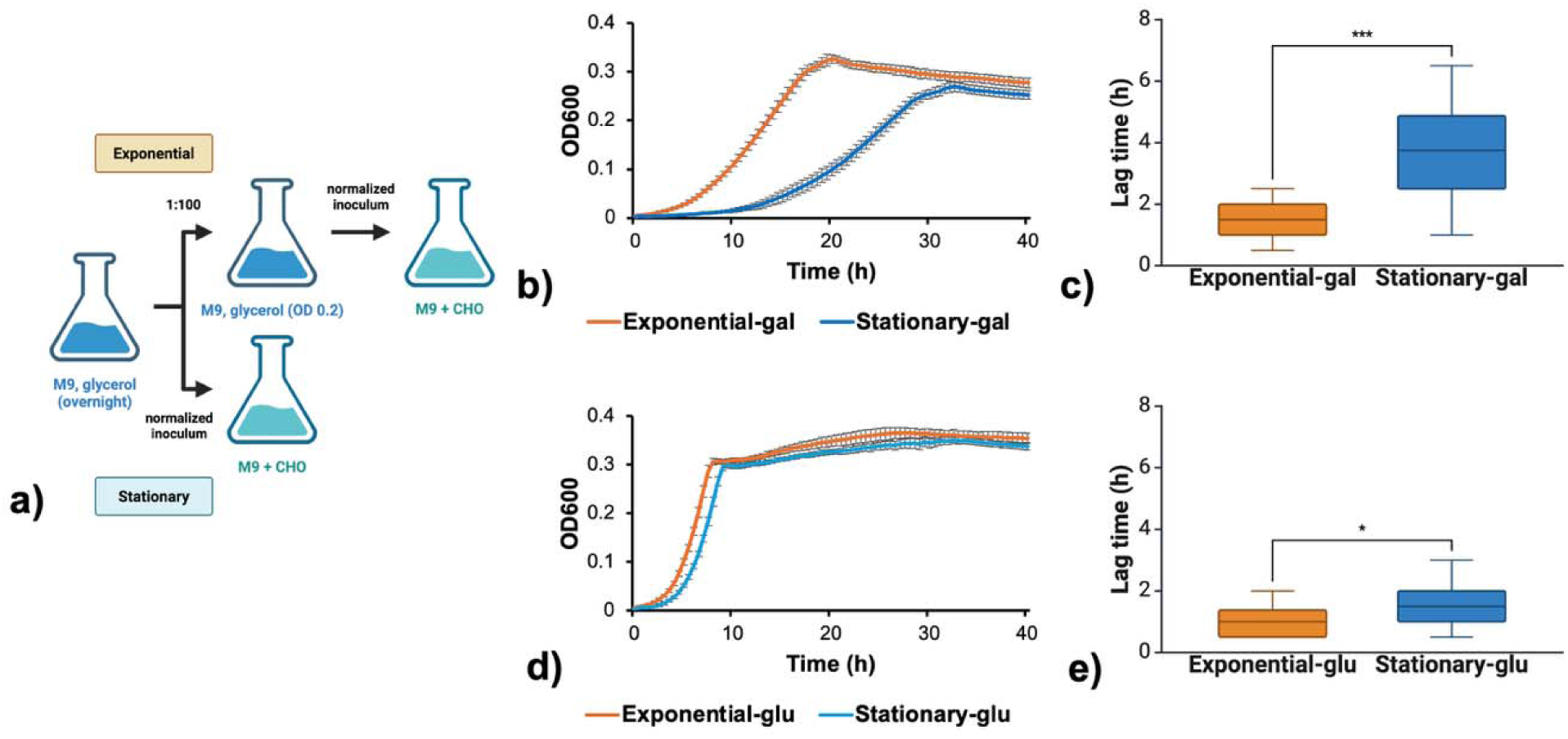
Exponential phase uniquely primes galactose metabolism. (a) Diagram of nutrient shift assay. (b, d) Growth curves) and (c, e) lag times of *E. coli* MG1655 cultures previously grown to exponential phase (OD_600_ 0.2) (orange) or stationary phase (blue) in M9+glycerol before dilution to the same density and transfer to M9+galactose (gal) or M9+glucose (glu). Data are represented as the mean ± standard deviation of three biological replicates, each of six technical replicates. Lag times were compared using a Welch’s t-test (* indicates p < 0.05, *** indicates p < 0.001).

Interestingly, cells passaged from exponential phase into M9+glucose exhibited only a slight growth advantage relative to those transferred from stationary phase (Fig. 3d-e). This finding further supported that the LB-to-M9 transition was costly, and that the growth enhancement following transfer to galactose was a unique HDB, which we will now refer to as “galactose HDB”.

### Galactose HDB is dependent on galactose repression

We hypothesized that the galactose repressors, GalR and GalS; as well as CRP, could influence galactose HDB, as they are the only known regulators of the *gal* operon. To test this, we passaged mutants with single- or double-gene deletions in *galR* and/or *galS* through exponential or stationary phase and tracked their growth following transfer to galactose. Here, HDB was lost in strains deficient in *galR* or *galS* (Fig. 4a-c, e). This was expected since deletion of either repressor should de-repress the *gal* operon and therefore enhance growth on galactose as a sole carbon source. Interestingly, deletion of both *galR* and *galS* did not improve growth on galactose further but caused a growth defect that reduced the endpoint density of the culture and increased lag time in a manner that was independent of passaging history (Fig. 4d-e). We could not test the effect of *crp*-deficiency on HDB, as a Δ*crp* strain would not grow in M9 supplemented with galactose as a sole carbon source (Fig. S1).

**Figure 4:**
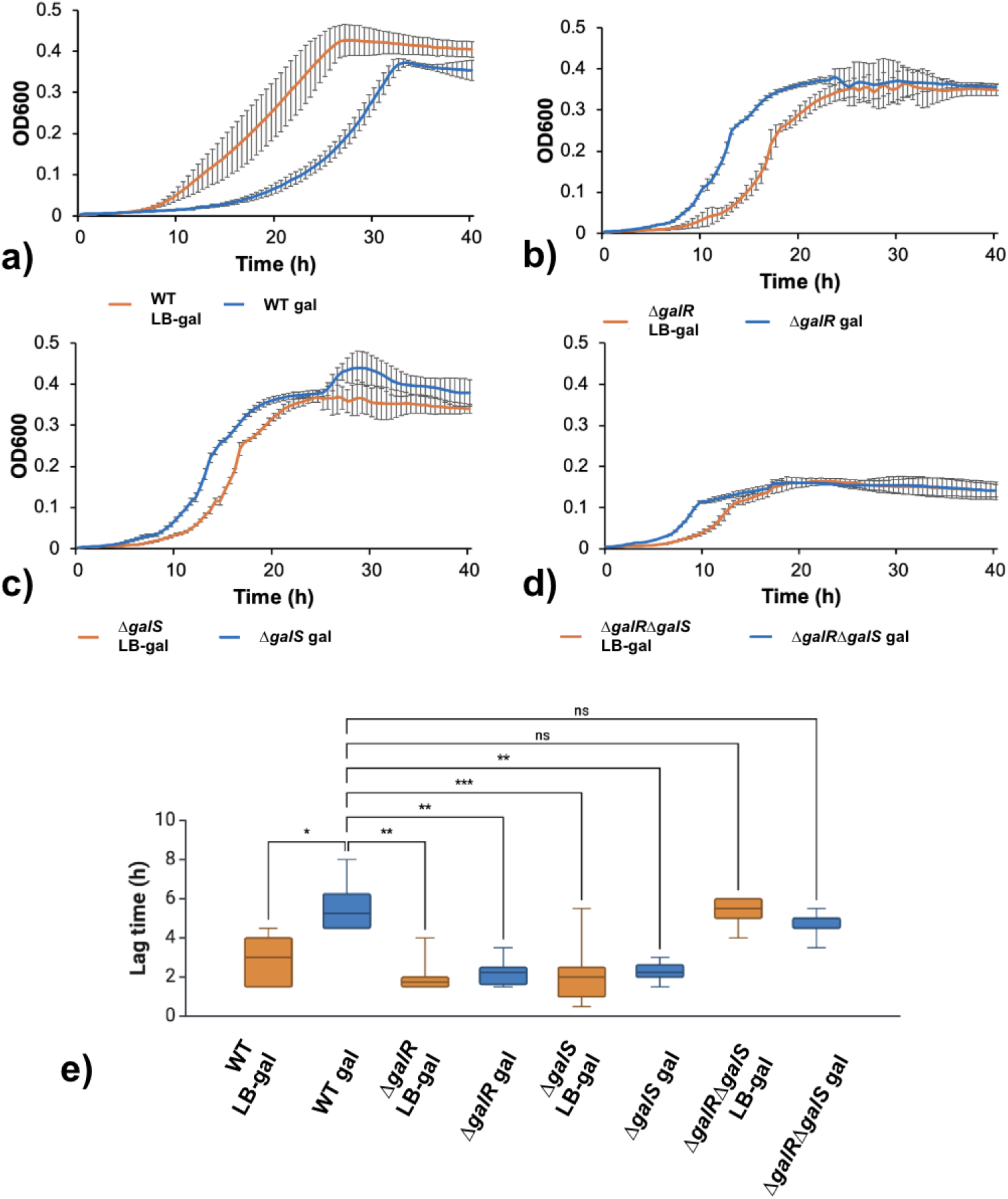
Primed galactose utilization is abrogated in repressor-deficient mutants. **(a-d)** Growth curves and (e) lag times of WT, Δ*galR*, Δ*galS*, and Δ*galR*Δ*galS E. coli* BW25113 previously grown to OD_600_ 0.2 in LB (orange) or stationary phase (blue) in M9+glycerol before diluted to the same density and transferred to M9+galactose. Data are represented as the mean ± standard deviation of three biological replicates, each of three technical replicates. Lag times were compared using a Kruskal-Wallis test with Dunn’s multiple comparisons test (* indicates p < 0.05, ** indicates p < 0.01, and *** indicates p < 0.001, “ns” indicates no statistical significance).

The loss of galactose HDB in repressor-deficient backgrounds suggested that the behaviour is a function of GalR and/or GalS. However, it was unclear how the repressors were tied to the HDB.

### Galactose HDB is maintained with increasing generations from stationary phase

To better understand the factors controlling the HDB, we passaged *E. coli* through M9+glycerol multiple times to increase the number of generations from stationary phase prior to transfer to M9+galactose (Fig. 5a). We reasoned that the galactose HDB could be related to intracellular repressor concentrations, which may accumulate during stationary phase but become diluted as cells resume growth. Alternatively, as described for *lac* HDB, daughter cells may inherit an activating factor, such as the galactose permease, that facilitates subsequent growth on galactose (Novick & Weiner, 1957). Here, we found that priming was maintained as the number of generations from stationary phase increased (Fig. 5b-c). This confirmed that HDB was tied to exponential growth, and that the HDB itself does not attenuate with further time or generations from stationary phase.

**Figure 5:**
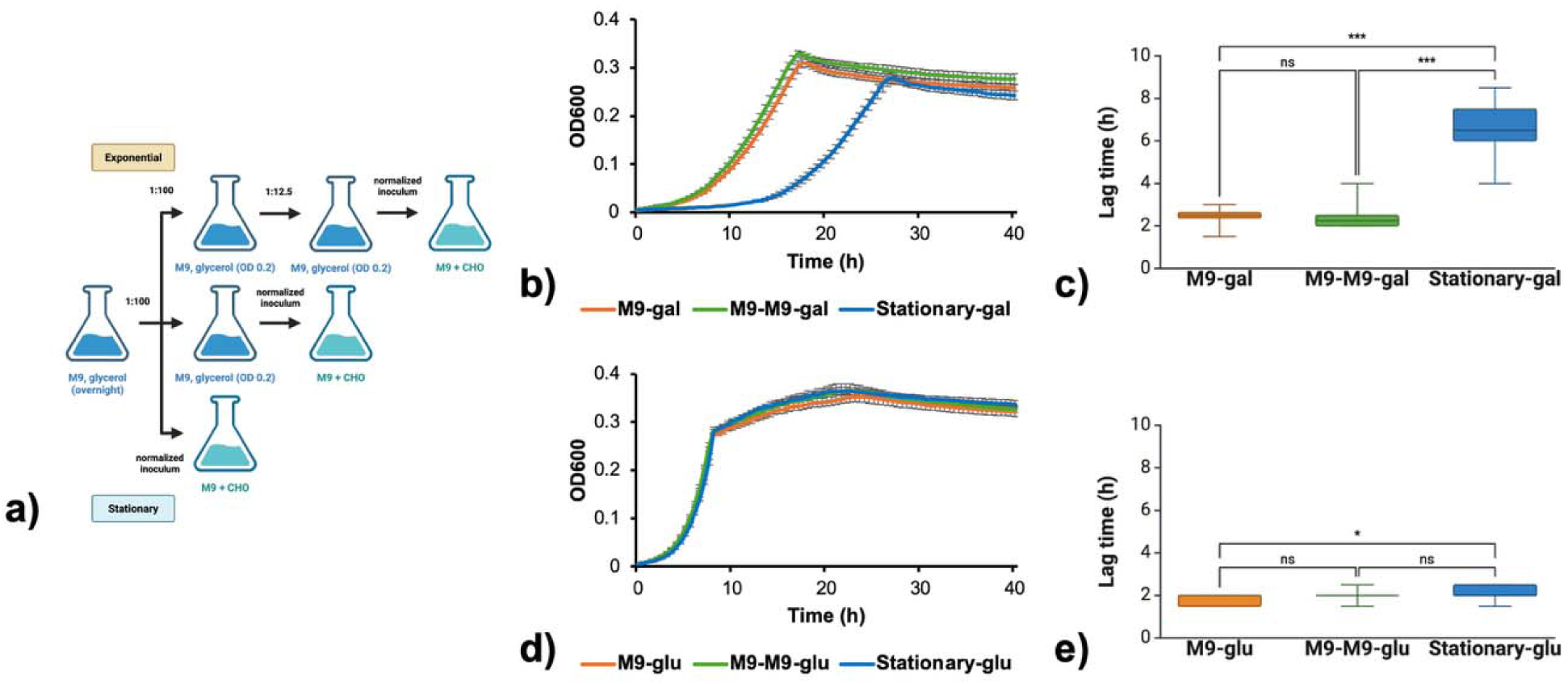
Primed galactose utilization is not influenced by increasing the number of generations from stationary phase in M9. (a) Diagram of nutrient shift assay. (b,d) Growth curves and (c, e) lag times of *E. coli* MG1655 cultures previously grown to OD_600_ 0.2 in M9+glycerol, or stationary phase (blue) in M9+glycerol before dilution to the same density and transfer to M9+galactose (gal) or M9+glucose (glu). The orange and green data were collected from cultures passaged through M9+glycerol once (M9) or two times (M9-M9), respectively. Note: the second M9+glycerol passage was performed using a lower dilution factor (1:12.5). Data are represented as the mean ± standard deviation of three biological replicates, each of four technical replicates. Lag times were compared using a Kruskal-Wallis test with Dunn’s multiple comparisons test (* indicates p < 0.05, *** indicates p < 0.001, “ns” indicates no statistical significance).

In contrast, cultures transferred into M9+glucose did not show any visual difference in their growth curves, regardless of whether they were passaged through exponential phase once or twice, or directly from stationary phase (Fig. 5d-e). A significant change in lag time was detected between cultures passaged directly from stationary phase into M9+glucose (stationary-glu) and those transferred after one exponential phase passage in M9+glycerol into M9+glucose (M9-glu). However, it is likely this effect was due to the limit of detection of the lag time quantification, as *E. coli* quickly transitions into exponential phase upon transfer to M9+glucose.

### Galactose HDB functions independently of GalR oligomerization

Learning that galactose metabolism could be primed in exponential phase suggested that the activity of GalR and/or GalS might be influenced by growth phase, as HDB was lost in both these repressor-deficient backgrounds. Previous work using chromosomal confirmation capture has shown that GalR dimers bind to distant operator sites and oligomerize to connect remote segments of DNA in 3D space (Qian et al., 2012). It has been proposed that these interactions help maintain a high local concentration of GalR around distant operators in the chromosome in stationary cells, as over 80% of these interactions disappear when cells are cultured to exponential phase (Qian et al., 2012). Considering that galactose HDB is tied to growth phase and is lost in a *galR*-deficient background, we hypothesized that the behaviour is a functional consequence of GalR-mediated chromosomal reorganization, by which repression of galactose metabolism is enhanced during stationary phase. To test this, we conducted a nutrient shift assay using a GalR mutant (GalR^T322R^) whichcan bind the *gal* operators but inefficiently forms tetramers or higher order oligomers that repress the operon through DNA looping (Agerschou et al., 2016; Geanacopoulos & Adhya, 2002). It has been shown through atomic force microscopy that on average, GalR^T322R^ forms fewer DNA loops than WT GalR, suggesting that it is defective in mediating intersegmental chromosomal connections (Csiszovszki et al., 2011; Qian et al., 2012). Therefore, galactose HDB should be lost in a GalR^T322R^ mutant if the behaviour is a result of GalR oligomerization and its associated effects on the chromosome. To our surprise, HDB was maintained, suggesting that the phenotype was not a function of GalR oligomerization (Fig. 6a-b).

**Figure 6:**
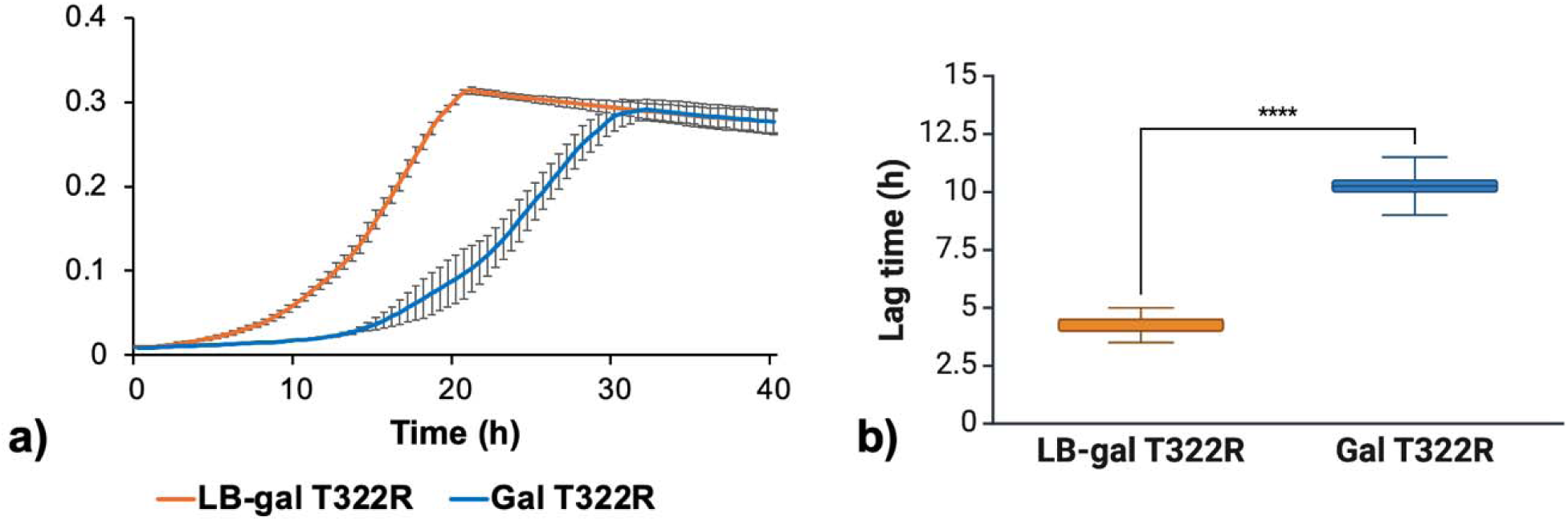
Primed galactose utilization is maintained in tetramerizing-deficient GalR^T322R^. **(a)** Growth curves and (b) lag times of *E. coli* MG1655 GalR^T322R^ previously grown to OD_600_ 0.2 in LB (LB-gal, orange) or stationary phase (Gal, blue) in M9+glycerol before dilution to the same density and transfer to M9+galactose. Data are represented as the mean ± standard deviation of three biological replicates, each of six technical replicates. Lag times were compared using a Welch’s t-test (**** indicates p < 0.0001).

### Galactose HDB is observed in diverse *E. coli* isolates

Given lab-adapted *E. coli* strains transition to exponential growth more frequently than microbes isolated from their natural environment, where resources are limited and constant bacterial growth is rarely found, se were curious whether galactose HDB was related to the natural history of *E. coli* (Navarro Llorens et al., 2010).

We assayed for galactose HDB in clinical *E. coli* isolates from fecal samples in an ulcerative colitis patient cohort. Interestingly, we found that galactose HDB was present in all strains tested, however, the magnitude of the behaviour varied across isolates, with some strains exhibiting a stronger phenotype than others (Fig. 7a-h). While the behaviour appeared to be generalizable to other *E. coli* strains, it was lost in *Salmonella* (Fig. 7i), one of the closest relatives in the *Enterobacteriaceae* family. This was surprising given that *E. coli* and *Salmonella* share the same genetic components and topographical structure of the *gal* regulon and also have a similar natural history as gut-associated bacteria (Csiszovszki et al., 2011).

**Figure 7:**
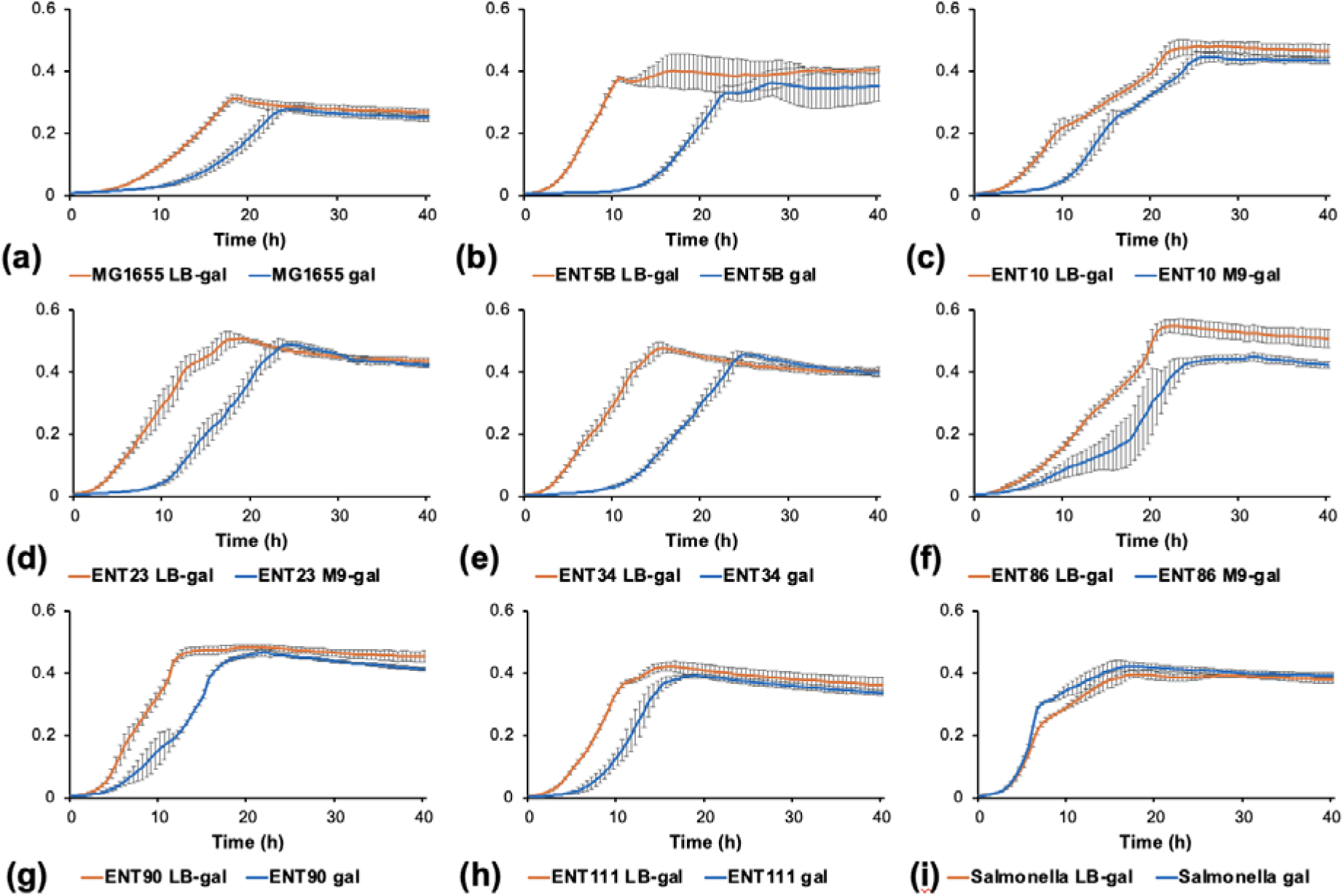
Primed galactose utilization is generalizable to *E. coli*, but not *Salmonella.* Growth curves of (a) *E. coli* MG1655, (b-h) clinical *E. coli* strains: ENT5B, ENT10, ENT23, ENT34, ENT86, ENT90, ENT111, and (i) *S. enterica* Typhimurium LT2 previously grown to OD_600_ 0.2 in LB (orange, LB-gal) or stationary phase (blue, gal) in M9+glycerol before dilution to the same density and transfer to M9+galactose. Data are represented as the mean ± standard deviation of three biological replicates, each of a minimum of six technical replicates.

## CONCLUSION

Here, we show evidence of a novel HDB in *E. coli* that links fitness in a galactose environment to the previous growth phase of the culture, wherein cells transferred to galactose media from exponential phase were primed for growth relative to cells transferred from stationary phase - and this was not true for other carbon sources. Although the complete mechanism underlying this HDB remains unclear, our findings implicate the galactose repressors (GalR and GalS) in mediating this behaviour. While it is tempting to suggest that this growth-phase-dependent response reflects an adaptive feature of *E. coli* physiology, perhaps shaped by recurring transitions between nutrient limitation and availability in the gut, the absence of the phenotype in *Salmonella* makes us wary of overinterpreting. Galactose metabolism is subject to multiple layers of regulation that distinguish it from the utilization of other carbon sources, including coordinated control by GalR/GalS, CRP-cAMP, and the galactose transport systems. Our findings expand the growing body of evidence that galactose metabolism is uniquely regulated among other carbohydrates, with cellular history shaping the response to a subsequent galactose environment. Together, our findings identify growth phase as a previously unrecognized determinant of galactose utilization in *E. coli* and provide a foundation for further investigation into the molecular mechanisms that encode and transmit this galactose-specific HDB.

## METHODS

### Bacterial strains and growth conditions

All strains were streak purified on lysogeny broth (LB) 1% (w/v) agar made with 1% (w/v) tryptone (Oxoid, Cat. # LP0042B), 0.5% (w/v) yeast extract (Fisher BioReagents, Cat. # BP1422-2), 1% (w/v) NaCl (VWR Chemicals, Cat. # BDH9286), and agar from Fisher BioReagents (Cat. # BP1423-500). Plates were incubated overnight at 37°C, from which single colonies were picked, inoculated into 5 mL LB, and incubated overnight at 37°C, shaking at 250 revolutions per minute (RPM) in the Infors HT Ecotron shaking incubator. Glycerol stocks were created by pelleting 1.5 mL of the overnight culture through centrifugation at 16,800 relative centrifugal force (RCF) for 1.5 min, and resuspending in 1 mL LB with15% glycerol (Fisher BioReagents, Cat. # BP299-4). Stocks were stored at −80°C. Strains were recovered by incubating a frozen scraping of glycerol stock with 5 mL of the desired medium at 37°C with agitation.

**Table 1:** Strains used in this study.

| Bacterium | Strain designation | Provenance |
| --- | --- | --- |
| *Escherichia coli* | MG1655 | (Blattner et al., 1997) |
| *Escherichia coli* | BW25113 | (Baba et al., 2006) |
| *Escherichia coli* | BW25113 $\Delta galR$ | (Baba et al., 2006) |
| *Escherichia coli* | BW25113 $\Delta galS$ | (Baba et al., 2006) |
| *Escherichia coli* | BW25113 $\Delta galS \Delta galR$ | This study |
| *Escherichia coli* | MG1655 GalR<sup>T322R</sup> | (Geanacopoulos & Adhya, 2002; Tao et al., 2023) |
| *Escherichia coli* | ENT5B | Surette Lab, McMaster (unpublished) |
| *Escherichia coli* | ENT10 | Surette Lab, McMaster (unpublished) |
| *Escherichia coli* | ENT23 | Surette Lab, McMaster (unpublished) |
| *Escherichia coli* | ENT34 | Surette Lab, McMaster (unpublished) |
| *Escherichia coli* | ENT86 | Surette Lab, McMaster (unpublished) |
| *Escherichia coli* | ENT90 | Surette Lab, McMaster<br>(unpublished) |
| *Escherichia coli* | ENT111 | Surette Lab, McMaster<br>(unpublished) |
| *Salmonella enterica* | Typhimurium LT2 | Félix d'Hérelle Reference Center<br>for Bacterial Viruses, Laval<br>(HER1023) |

### Nutrient shift assay

A frozen scraping from a glycerol stock of the desired bacterium was used to inoculate 5 mL autoclaved M9 minimal medium supplemented with the following filter-sterilized reagents: 2 mM MgSO_4_, 0.1 mM CaCl_2_, and 0.4% glycerol. The culture was incubated at 37°C overnight, with agitation. The next day, the overnight culture was sub-inoculated by creating a 1:100 dilution into 5 mL of the conditioning medium (LB or M9 supplemented with a carbon source). The subculture was grown to early exponential phase (OD_600_ 0.2) under the same conditions as the overnight culture. OD_600_ was measured using the Spectronic 20D+ spectrophotometer (Thermo Scientific, MA, USA). The overnight and subculture were pelleted through centrifugation at 7000 RCF for 5 min, washed with 5 mL M9 without a carbon source, and resuspended in M9 to a normalized density between OD_600_ 0.1-0.2. The resuspended cultures were added to a 96-well plate containing M9 supplemented with an experimentally pre-determined limiting concentration of a carbon source (5 mM for monosaccharides and 2.5 mM for disaccharides). All carbon sources were filter-sterilized and obtained from Thermo Fisher Scientific (MA, USA). Growth was monitored every 30 mins for 18-40 h in an Epoch 2 microplate spectrophotometer heated to 37°C with double orbital agitation. Note: the Synergy H1 Multimode microplate reader (BioTek Instruments, VT, USA) was used occasionally depending on microplate reader availability. Wells containing media alone served as a control for media contamination. Data were represented as a growth curve normalized to the background absorbance of the growth medium.

Lag time was estimated using a modified version of the biomass increase method, which measures the time required for the population size to increase by a certain value from the beginning of growth (Opalek et al., 2022; Smug et al., 2024). However, instead of applying a fixed threshold, lag time was quantified as the amount of time required for the initial population to double in optical density. This approach allowed for more consistent comparisons across conditions with varying initial densities.

### FLP-FRT recombination and P1 transduction

A double knockout of the two galactose repressors (*galR* and *galS*) was created by moving the Δ*galR* deletion into a kanamycin-sensitive Δ*galS* strain of *E. coli* BW25113 according to the P1 transduction protocol described by (Deatherage et al., 2021). The kanamycin-sensitive Δ*galS* recipient was generated through FLP-FRT recombination following the protocol described by (Barrick et al., 2023). The pCP20 vector for FLP-FRT recombination and P1 lysate for the transduction were graciously provided by the Brown Lab at McMaster University. The genotype of the resulting strain was confirmed by PCR using primers flanking each of the *galR* and *galS* genes. Each PCR was set up as a 25 μL reaction containing 1 μL of each forward and reverse primer, 2.5 μL 10x PCR buffer, 0.5 μL dNTPs, 0.25 μL Taq DNA polymerase, and 1 μL of an overnight culture grown in LB. The remaining volume consisted of nuclease free water. Primers were ordered from Integrated DNA Technologies (IA, USA) and all other PCR reagentswere obtained from FroggaBio (Cat. #T-500). Thermal cycling consisted of an initial denaturation for 5 min at 95 °C followed by 35 cycles of the following steps: denaturation at 94 °C for 45 s, annealing at 60 °C for 45 s, and extension at 72 °C for 1.5 min. A final extension was performed at 72 °C for 5 min. PCRs were performed in duplicate with controls containing cultures of WT, and single-gene deletion Δ*galR* and Δ*galS E. coli* BW25113 cultures.

**Table 2:**
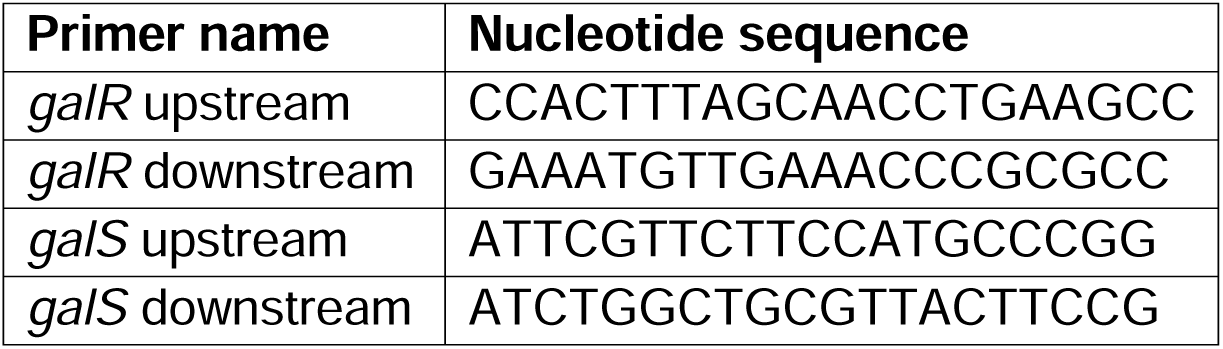
Oligonucleotides.

## ACKNOWLEDGEMENTS

We thank Sankar Adhya’s laboratory at the National Cancer Institute (NCI), Center for Cancer Research, Laboratory of Molecular Biology, for providing the *E. coli* GalR^T322R^ mutant *E. coli* strain (MD, USA). We also thank Dominique Tertigas of Michael Surette’s laboratory at McMaster University for providing clinical *E. coli* isolates from an ulcerative colitis patient cohort.

## FUNDING

This work was supported by funding from the Weston Family Foundation and the Natural Sciences and Engineering Research Council of Canada (NSERC). APH is supported by the Farncombe Family Chair in Phage Biology. MGS …

## SUPPLEMENTARY MATERIAL

**Figure S1:**
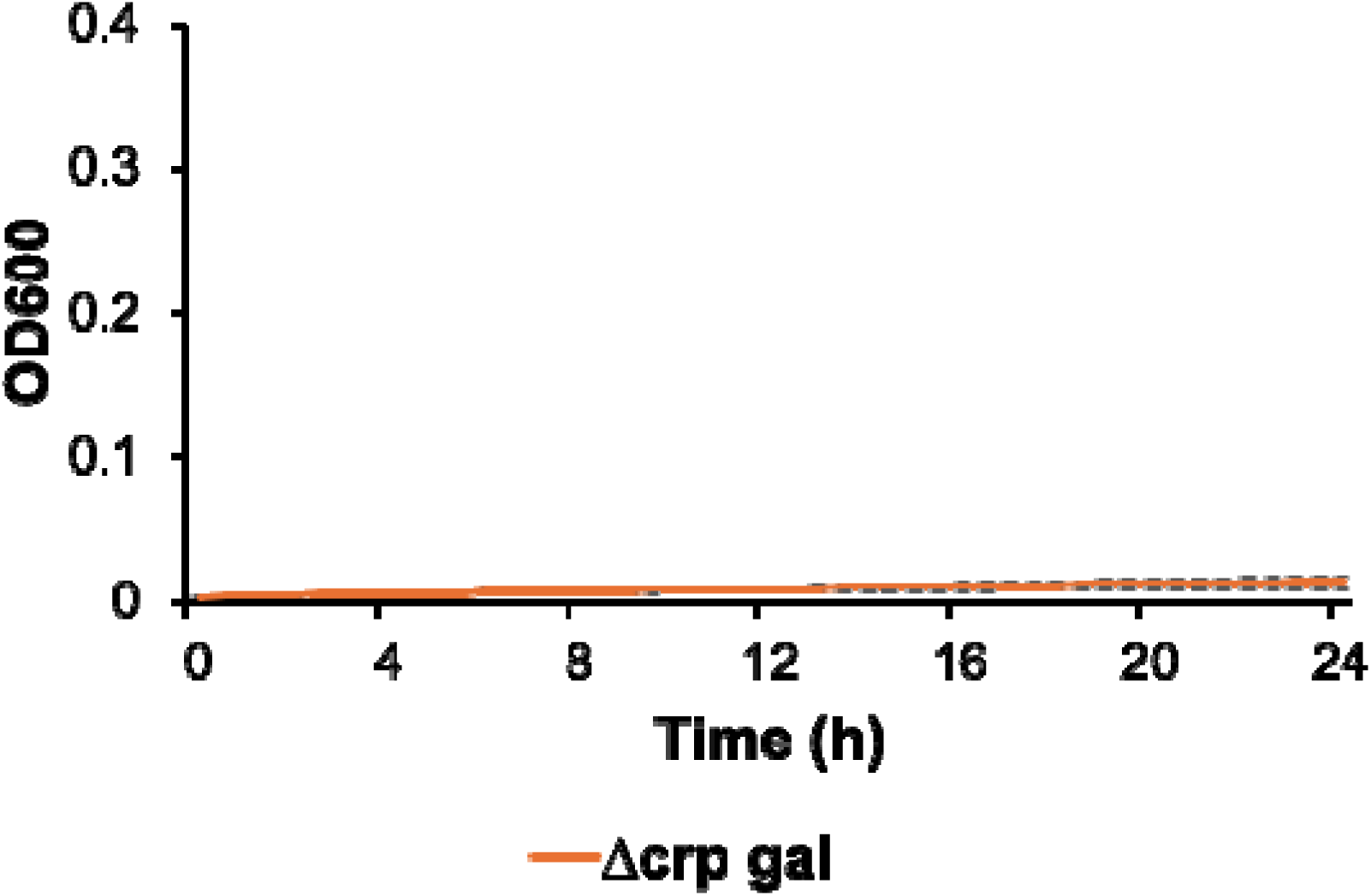
Growth of *E. coli* Δ*crp* on M9+galactose. Growth curves of *E. coli* MG1655 Δ*crp* previously grown on M9+galactose. Data are represented as the mean ± standard deviation of three biological replicates, each of six technical replicates.

## Notes

### Competing Interest Statement

The authors have declared no competing interest.

